# The metabolically protective energy expenditure increase of *Pik3r1*-related insulin resistance is not explained by Ucp1-mediated thermogenesis

**DOI:** 10.1101/2024.02.12.579851

**Authors:** Ineke Luijten, Ami Onishi, Eleanor J. McKay, Tore Bengtsson, Robert K. Semple

**Affiliations:** Centre for Cardiovascular Science, University of Edinburgh, Edinburgh, UK; Department of Molecular Biosciences, The Wenner-Gren Institute, Stockholm University, Stockholm, Sweden; MRC Human Genetics Unit, Institute of Genetics and Cancer, University of Edinburgh, Edinburgh, UK

**Author notes:** **Correspondence to:** Prof. Robert K. Semple, Centre for Cardiovascular Science, University of Edinburgh, Queen’s Medical Research Institute, Little France Crescent, Edinburgh EH16 4TJ, UK. /.

**Keywords:** Insulin resistance, PI 3-Kinase, Pik3r1, energy expenditure, brown adipose tissue

## Abstract

Human SHORT syndrome is caused by dominant negative human *PIK3R1* mutations that impair insulin-stimulated phosphoinositide 3-kinase (PI3K) activity. This produces severe insulin resistance (IR) and often reduced adiposity, commonly described as lipodystrophy. However unlike human primary lipodystrophies, SHORT syndrome does not feature fatty liver or dyslipidaemia. *Pik3r1^Y657*/WT^* (Pik3r1^Y657*^) mice metabolically phenocopy humans, moreover exhibiting increased energy expenditure. We have hypothesised that this increased energy expenditure explains protection from lipotoxicity, and suggested that understanding its mechanism may offer novel approaches to mitigating the metabolic syndrome. We thus set out to determine whether increased Ucp1-dependent thermogenesis explains the increased energy expenditure in *Pik3r1*-related IR. Male and female Pik3r1^Y657*^ mice challenged with a 45% fat diet for 3 weeks at 21°C showed reduced metabolic efficiency not explained by changes in food intake or physical activity. No changes were seen in thermoregulation, assessed by thermal imaging and a modified Scholander protocol. Ucp1-dependent thermogenesis, assessed by norepinephrine-induced oxygen consumption, was also unaltered. Housing at 30°C did not alter the metabolic phenotype of male Pik3r1^Y657*^ mice, but led to lowered physical activity in female Pik3r1^Y657*^ mice compared to controls. Nevertheless these mice still exhibited increased energy expenditure. Ucp1-dependent thermogenic capacity at 30°C was similar in Pik3r1^Y657*^ and WT mice. We conclude that the likely metabolically protective ‘energy leak’ in Pik3r1-related IR is not caused by Ucp1-mediated BAT hyperactivation, nor impaired thermal insulation. Further metabolic studies are required to seek alternative explanations such as non Ucp1-mediated futile cycling.

**New and Noteworthy:** Understanding how Pik3r1^Y657*^ mice and humans are protected from lipotoxicity despite insulin resistance may suggest new ways to mitigate metabolic syndrome. We find reduced metabolic efficiency and increased energy expenditure in Pik3r1^Y657*^ mice but no differences in locomotion, thermoregulation or Ucp1-dependent thermogenesis. Protective energy expenditure in Pik3r1-related insulin resistance has an alternative, likely metabolic, explanation

## Introduction

Insulin resistance (IR) is generally associated with dyslipidaemia, characterised by high serum triglyceride, and low HDL cholesterol, and associated with fatty liver (e.g. [1,2]). This is also true in most forms of severe IR of known aetiology [3,4]. We and others have reported, however, that monogenic severe IR due to loss of function of the insulin receptor, or downstream phosphoinositide 3-kinase (PI3K), are an exception to this. Specifically, in SHORT syndrome, which features severe IR and is caused by heterozygous *PIK3R1* mutations, serum triglyceride concentrations and liver triglyceride contents are surprisingly normal or low [5]. These findings demonstrate that IR is uncoupled from dyslipidaemia in humans when caused by primary proximal insulin signalling defects. What explains this beneficial IR subphenotype is unknown, and elucidating the mechanism holds the promise of uncovering new strategies to mitigate IR-related comorbidities.

We have described a mouse model of SHORT syndrome caused by the *Pikr3r1* Y657* mutation [6]. This mutation truncates all three protein products of the *Pik3r1* gene - p85α/p50α/p55α - in the C-terminal SH2 domain [7–10]. In keeping with this, mice heterozygous for *Pik3r1* Y657* (*Pik3r1^Y657*^* mice) show attenuated intracellular insulin signalling, and thus severe IR [5,6,11]. *Pik3r1^Y657*^* mice also show reduced adiposity, a defining feature of lipodystrophy, but while lipodystrophy is associated with severe dyslipidaemia and fatty liver, neither increased liver fat, nor hyperlipidaemia are seen in *Pik3r1^Y657*^* mice compared to wild-type (WT) littermates [6], phenocopying the human IR phenotype caused by PI3K hypofunction. This murine model is thus valid for interrogation of the mechanism underlying protection from dyslipidaemia despite IR.

Studies to date have demonstrated increased energy expenditure in *Pik3r1^Y657*^* mice [6], and indeed in several other models of pharmacologically or genetically impaired PI3K signalling [12–19]. This increased energy expenditure could explain normolipidaemia despite IR in SHORT syndrome. Due to the lipid-lowering effect of brown adipose tissue (BAT) activation in hyperlipidaemic mice [20], it may be that BAT, and to a lesser extent white adipose tissue (WAT) thermogenesis explains the energy expenditure increase caused by PI3K inhibition. This hypothesis has been advanced previously, but evidence presented to date has not been definitive, and no mechanism whereby impaired PI3K signalling may activate BAT thermogenesis has been proven [12–14,17,18,21]. Moreover, any BAT activation observed may be secondary to changes in insulation or heat loss in *Pik3r1^Y657*^* mice, as is shown in mouse models of other pathologies [22,23]. This is particularly relevant as all studies in mouse models of impaired PI3K signalling have been performed below thermoneutrality (i.e. at 21°C), when BAT thermogenesis is mainly determined by sympathetic drive and animals’ cold perception [24].

We now set out to investigate the source of the unexplained “energy leak” of the *Pik3r1* Y657* mouse model of SHORT syndrome, assessing BAT activity, heat loss through the tail vein, and thermal insulation.

## Materials and methods

### Animal model and maintenance

Generation of mice carrying the *Pik3r1* Y657* mutation and an intronic neomycin resistance cassette flanked by loxP sites was previously described [6]. For this study the selection cassette was excised by crossing these mice with CAG-cre expressing mice. Mice heterozygous for the *Pik3r1* Y657* mutation (hereafter *Pik3r1^Y657*^*mice) and WT littermate controls were derived by breeding *Pik3r1^WT/WT^*and *Pik3r1^WT/Y657*^* mice on a C57Bl6/J background. All genotyping was performed by Transnetyx. No experimental animals expressed cre. Animals were group-housed in individually ventilated cages either at the Biological Research Facility at the University of Edinburgh or at the Experimental Core Facility at Stockholm University, both of which maintained a 12 h/12 h light/dark cycle and a controlled standard humidity (50%). Experiments were carried out under the UK Home Office Animals (Scientific Procedures) Act 1986, following University of Edinburgh ethical review, or under approval by the North Stockholm Animal Ethics Committee.

Until 10 weeks of age, mice had free access to water and chow diet (CRM, Special Diets Services at University of Edinburgh; Altromin 1324P, Brogaarden at Stockholm University), and were housed at 21°C. At 10 weeks of age, mice were single-caged and either kept at 21°C or moved to 30°C. Experimental procedures began after a 2-week acclimatization period, when all animals were put on a 45% high-fat diet (HFD, D12451, Research Diets) provided *ad libitum*. Animals received HFD for a maximum of 3 weeks, during which the experimental procedures outlined below were performed in various cohorts.

At the end of the experimental period, animals were culled by exposure to increasing concentrations of CO_2_, except after the norepinephrine test (see 2.6), when animals were already sedated by pentobarbital injection. When unconscious, nose-anus length was measured using a digital calliper, after which death was confirmed by cervical dislocation. Both lobes of interscapular brown adipose tissue (BAT) and inguinal white adipose tissue (ingWAT) were collected immediately post-mortem, weighed, snap-frozen in liquid nitrogen, and stored at - 80°C. For determination of tibia length, the left leg was put in 30% KOH for 45 min, after which the tibia was isolated and its length measured with a digital calliper.

### Determination of metabolic efficiency

Body weight and food weight were measured weekly from 11 weeks of age. Food intake in kJ/day was calculated by dividing weekly food intake in grams by 7 and multiplying by the energy density of the diets (Chow CRM 10.74 kJ/g; Chow Altromin 13.5 kJ/g; HFD 19.8 kJ/g). Body composition was measured by tdNMR using the Bruker Minispec Live Mice Analyzer LF50 (University of Edinburgh), or by MRI using the EchoMRI-100^TM^ (Echo Medical Systems, USA. Stockholm University) at the start (12 weeks old) and end (14 weeks old) of the HFD-feeding period. Change in body composition was calculated as (fat mass end– fat mass start) + (lean mass end– lean mass start) and converted to kJ using 39 kJ/g and 5 kJ/g as energy densities for fat and lean mass respectively. Metabolic efficiency was calculated by dividing the difference in body composition (in kJ) by food intake (in kJ) over the same period, multiplied by 100.

### Characterisation of energy metabolism

After 2 weeks of HFD feeding (age 14 weeks), animals were placed in temperature-controlled metabolic chambers (Promethion Core system, Sable Systems) for 1 week, using random assignment to cages and analysers. In the Promethion system, mice were housed at the same temperature that they had been housed at throughout the study (either 21°C or 30°C), with a 12 h/12 h light/dark cycle and free access to HFD and water. O_2_ consumption, CO_2_ production, food intake, water intake, and physical activity (XYZ beam breaks) measurements were collected in three-minute intervals. Data collected during the first 3 days were excluded from analyses, as this was considered an acclimatisation period. Energy expenditure during days 4-6 was calculated in watts using a modified Weir equation:

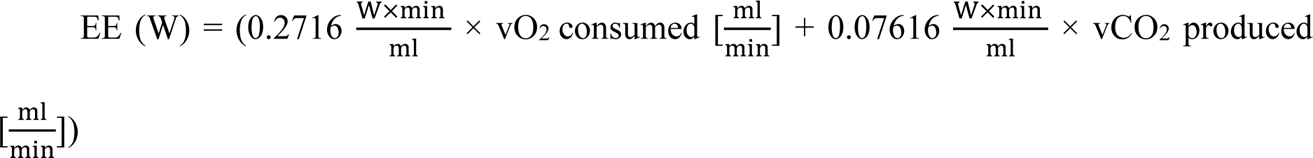

[23,25].

Food intake was set to 0 at 7AM on day 4 and calculated cumulatively thereafter, before conversion from g to kJ using the energy density of the HFD (see 2.2). Physical activity was calculated as the sum of X-, Y- and Z-bream breaks per time point. The theoretical thermic effect of food was calculated using the thermic effects of fat (2.5%), carbohydrates (7.5%) and protein (25%) [26].

### Assessment of body temperature and heat loss

After 1 week of HFD-feeding (age 13 weeks), tail and inner ear temperatures were measured using a FLIR^®^ T650sc infrared camera (FLIR Instruments, Germany). To prevent warming of the tail by touching, animals were moved in a cardboard tunnel and placed on top of their cage lid, where they could move freely. At least 4 images were taken per animal from the inner ear and 4 from the tail. Inner ear, tail base and tail temperatures were determined for each image using FLIR Tools Software.

### Assessment of thermal insulation

To measure thermal insulation, a modified Scholander experiment [27] was performed. After 2 weeks of HFD-feeding, animals (age 14 weeks) were placed in the Promethion system at 21°C with free access to HFD and water. After a 2.5-day acclimatisation period, the system temperature was increased to 33°C, maintained for 2h, and then decreased sequentially to 30°C, 25°C, 20°C and 15°C, with every temperature maintained for 2h. O_2_ consumption, CO_2_ production and physical activity measurements were collected and analyzed. O_2_ consumption values at each temperature were averaged from 5 successive points when physical activity, measured by XYZ-beam breaks, was closest to zero. Energy expenditure per gram of lean mass was calculated as described in 2.3.

### Determination of Ucp1-dependent thermogenic capacity

To determine total Ucp1-dependent thermogenic capacity, a norepinephrine (NE) test was performed. Animals were sedated by intraperitoneal (i.p.) injection of 90 mg/kg pentobarbital (APL, Sweden), and placed in individual glass cylinders in the Promethion system at 30 °C. Baseline O_2_ consumption was measured in three-minute intervals for 20 minutes. Animals were briefly taken out of the cylinders and injected i.p. with 1 mg/kg norepinephrine bitartrate salt monohydrate (Sigma, Germany). Animals were returned to the cylinders and O_2_ consumption was measured for 40 minutes. Data was analyzed with O_2_ consumption expressed per gram lean mass. Total Ucp1-dependent thermogenic capacity was calculated by subtracting baseline O_2_ consumption (average of 3 successive stable points) from peak NE-induced O_2_ consumption (average of 3 successive highest points).

### Western blot analyses

Total Ucp1 protein in BAT and ingWAT lobes was determined by western blot analysis, detailed methods can be found in online supplementary materials. Adipose tissues were homogenized in RIPA buffer containing protease (cOmplete^TM^ Mini, Roche, Germany) and phosphatase (5 mM Na fluoride, 1 mM Na orthovanadate) inhibitors. Proteins were separated on a 12% polyacrylamide gel (Mini-PROTEAN® TGX^TM^, BioRad), and transferred to a PVDF membrane (BioRad) using a Transblot Turbo system (BioRad) according to manufacturers’ instructions. Membranes were blocked in 5% milk (w/v; Semper) in TBS-T and probed with primary anti-Ucp1 antibody (1:15000 solution in 5% milk in TBS-T, gifted by Jan Nedergaard and Barbara Cannon) and anti-αTubulin (1:1000 solution in 5% milk in TBS-T, Abcam 2144). After incubation with secondary anti-rabbit antibody (Cell Signaling 7074, 1:10000 in 5 % milk in TBS-T for Ucp1 measurements, 1:1000 for αTubulin measurements). proteins were visualized using Clarity^TM^ Western ECL Substrate (BioRad) and imaged with a ChemiDoc^TM^ XRS+ Molecular Imager (BioRad). The obtained images were analyzed with Image Lab^TM^ software (BioRad).

### Statistical analysis

All statistical analyses were undertaken in Graphpad Prism, version 10.1.1. No formal a priori power calculations were undertaken, but sample sizes drew on effect sizes for energy expenditure observed in prior studies of *Pik3r1^Y657*^* mice [6]. The only exclusion criteria related to predefined wellbeing endpoints, and these were not reached in any experimental group. Normality of data was confirmed using Kolmogorov-Smirnov testing. All data are represented as mean ± SEM. For comparison of differences between two groups at a single time-point, a Student’s t-test was used. Differences in one parameter between two groups over time were examined by two-way ANOVA followed by Šídák’s multiple comparisons test. For comparison of energy expenditure, cumulative food intake, and beam break data, the area under the curve (AUC) was determined per animal. Averages and standard errors were calculated per genotype and compared by Student’s t-test. For comparison of slopes and Y-intercepts of energy expenditure in relation to ambient temperature between genotypes, linear regression was performed on data points ranging from 15°C to 30°C. Linear regression was also used to assess energy expenditure in relation to lean body mass, with “p” values indicated relating to testing of differences in y interceptors of regression lines. All study data are reported.

## Results

### Pik3r1^Y657*^ mice have decreased metabolic efficiency

We previously reported that heterozygous *Pik3r1^Y657*^*exhibit increased energy expenditure after 8 weeks’ of high fat feeding that is not explained by increased locomotor activity [6]. We first set out to reproduce this observation, now studying mice in which the intronic antibiotic resistance cassette had been excised, conducting studies in two different animal facilities, and using a shorter term dietary intervention.

Individually housed 12-week old WT and *Pik3r1^Y657*^* mice were challenged calorically with a 45% HFD, having been chow fed to that point. This approach is commonly used to impose a load on adipose tissue and unmask lipodystrophic phenotypes [6]. As previously reported [6], both male and female *Pik3r1^Y657*^* mice were smaller than WT littermates (Fig. 1A), with lower body weight, body length, and tibia length (Fig. 1A-D). The reduced body weight in *Pik3r1^Y657*^* mice was accounted for by reduction of both lean and fat mass (Figs. 1E,F). Despite their smaller size, *Pik3r1^Y657*^* mice tended to eat more than WT mice (Fig. 1G baseline). During the first two weeks of HFD feeding, both male and female WT mice became hyperphagic, an effect that decreased by 3 weeks in males and plateaued in females (Fig. 1G). In contrast, *Pik3r1^Y657*^* mice showed a smaller increase in food intake from their elevated baseline, and this increase was sustained for both males and females.

**Figure 1.**
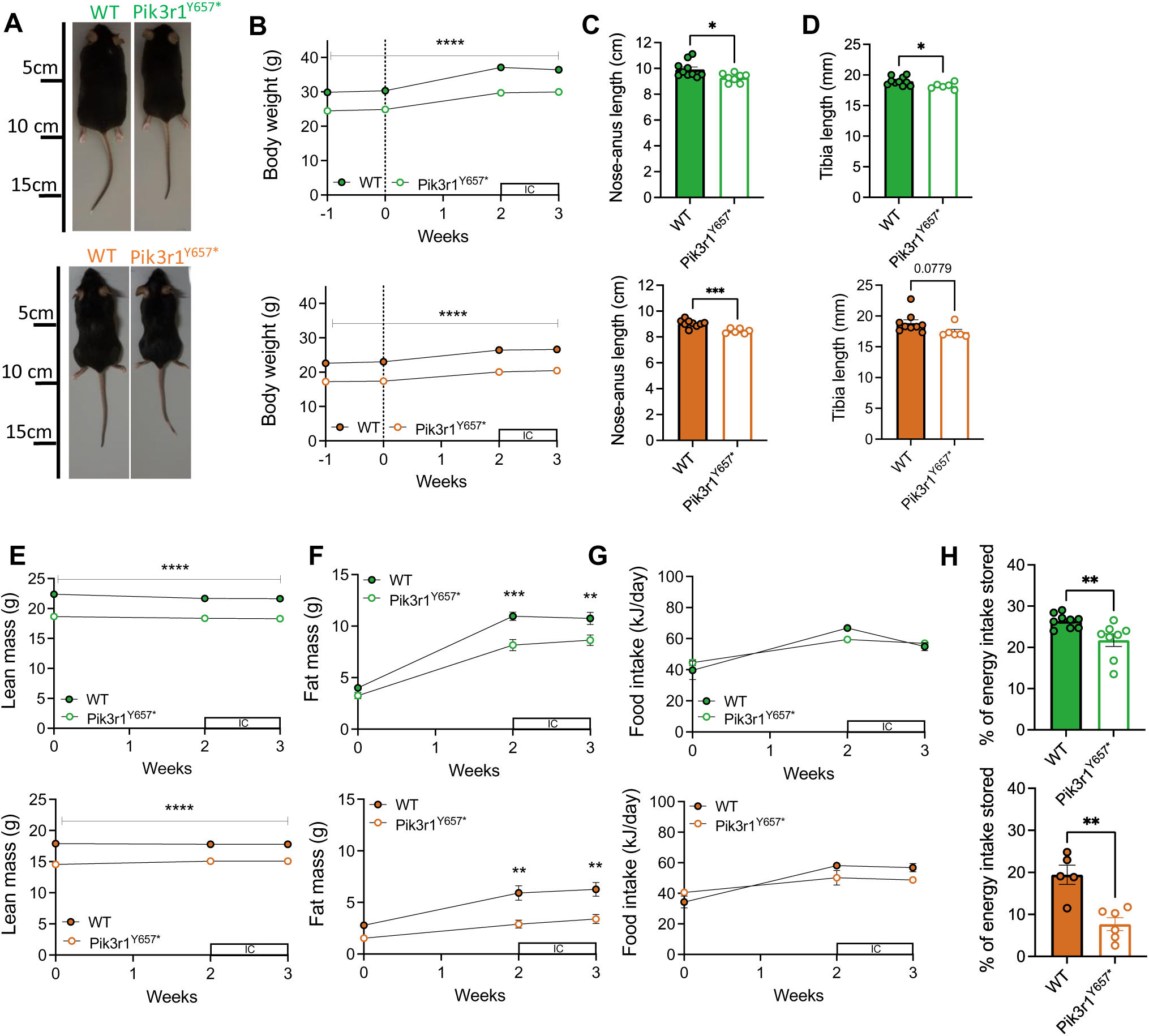
Male and female *Pik3r1^Y657*^* mice at 21°C show reduced metabolic efficiency. Male (green; top images) and female (orange; bottom images) WT and *Pik3r1^Y657*^* mice were fed a 45% HFD from t=0 (12 weeks old, indicated by dotted lines) and kept at 21°C throughout the study. Boxes with ‘IC’ represent periods in the indirect calorimetry system. (A) Representative images at 15 weeks’ old. (B) Body weight over time. (C) Body length (nose-anus) at 15 weeks old. (D) Tibial length at 15 weeks’ old. (E) Lean mass over time. (F) Fat mass over time. (G) Food intake over time. (H) Metabolic efficiency calculated between 12-14 weeks’ old. All data are represented as mean ± SEM; error bars may be too small to be visible; Male WT N=10, Male *Pik3r1^Y657*^* N=8, Female WT N=9, Female *Pik3r1^Y657*^* N=6. * P<0.05, ** p<0.01, *** p<0.001 and **** p<0.0001. Two-way ANOVA with Šídák’s multiple comparisons test, significance per timepoint between genotypes (B, E-G); Student’s t test (C, D, H).

Based on changes in lean and fat mass and the total food intake over the first two weeks of HFD feeding, we next calculated metabolic efficiency. Both male and female *Pik3r1^Y657*^* mice stored less of the energy they consumed than WT littermates (Fig. 1H). As we have previously found no evidence for nutritional malabsorption in mice of this genotype [6], this suggests increased energy expenditure as the likely explanation for reduced fat tissue accretion.

### Increased energy expenditure in Pik3r1^Y657*^ mice is not explained by increased food intake or physical activity

To confirm that decreased metabolic efficiency of *Pik3r1^Y657*^* mice was indeed associated with greater energy expenditure compared to WT mice, as indicated by our prior longer term studies [6], we next undertook indirect calorimetry during the third week of HFD-feeding. In neither male nor female *Pik3r1^Y657*^* mice was there any significant difference in total energy expenditure compared to wildtype littermates, however when lean mass was taken into account relatively increased energy expenditure was seen in male but not female *Pik3r1^Y657*^* mice (Figs. 2A, E).

**Figure 2.**
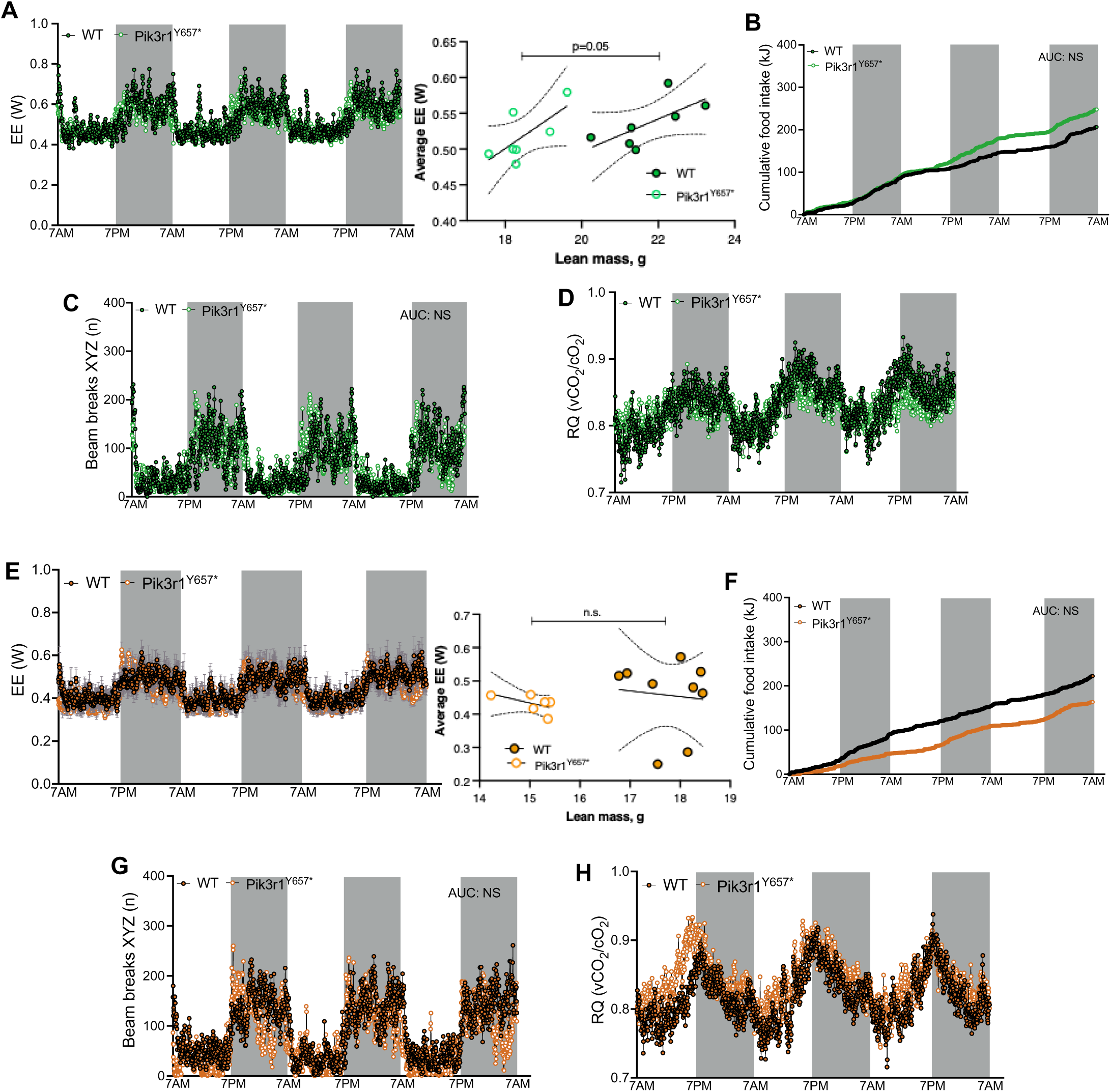
Energy metabolism in *Pik3r1^Y657*^* mice at 21°C. Male (green) and female (orange) WT and *Pik3r1^Y657*^* mice were fed a 45% HFD and kept at 21°C throughout the study. Grey boxes represent dark phases. (A,E) 72h energy expenditure. Raw traces are shown with adjacent linear regression plots of the last 24 hours’ energy expenditure against lean mass. Regression lines and 95% confidence interval are shown, with p value relating to testing of differences in intercept (no difference in gradients) (B, F) Cumulative food intake calculated in kJ. (C, G) Sum of X, Y and Z beam breaks per time point. (D,H) Respiratory quotient. All data are represented as mean ± SEM (shaded areas); Male WT N=10, Male *Pik3r1^Y657*^* N=8, Female WT N=9, Female *Pik3r1^Y657*^* N=6. NS = non-significant, **** p<0.0001. AUC determined per animal and averaged per genotype, compared by Student’s t test (A-F).

To investigate the origin of the increased energy expenditure, we assessed whether there were any changes in the obligatory components of energy expenditure, i.e. standard metabolic rate, diet-induced thermogenesis, and physical activity associated with activities such as feeding [28]. No significant differences in food intake were found between genotypes for either sex, with trends observed in different direction for each (Figs 1B,F), indicating that increased energy expenditure seen in *Pik3r1^Y657*^* mice of both sexes cannot be explained by differences in obligatory diet-induced thermogenesis.

Next, we counted XYZ-beam breaks as a combined index of obligatory (feeding), and facultative (e.g. exercise and fidgeting) physical activity. No changes in physical activity between male or female *Pik3r1^Y657*^*mice and WT littermates (Figs. 2C, G), excluding this potential cause for increased energy expenditure. No consistent difference in respiratory exchange ratio was observed between genotypes across sexes (Figure 2D,H).

### No change in thermoregulation and BAT activity in Pik3r1^Y657*^ mice

We next assessed cold-induced thermogenesis, mediated by skeletal muscle (shivering thermogenesis) and/or BAT (non-shivering thermogenesis) [28]. Because all mice were housed at a temperature well below thermoneutrality (i.e. standard ambient animal house temperature of 21°C [24]) throughout the study, it was plausible that any difference in cold perception between WT and *Pik3r1^Y657*^* mice could have translated into a difference in thermoregulatory response, and thus increased energy expenditure, in order to defend body temperature.

To characterize thermoregulation of *Pik3r1^Y657*^*mice, we first measured body temperature from the inner ear of the mice using an infrared camera. We found no difference between genotypes in the males, but a small decrease in female *Pik3r1^Y657*^* mice compared to WT (Fig. 3A). We next determined whether the mice showed any differences in heat dissipation through the tail vein, an important mechanism whereby mice defend their body temperature at high (via vasodilation) or low (via vasoconstriction) temperatures, which could plausibly be affected by altered PI3K signaling [29–31]. Again using an infrared camera, we measured tail temperatures at both base and the middle section (Fig. 3B), but found no differences between genotypes in either males or females (Figs. 3C-D), indicating no detectable difference in tail vasoregulation and heat loss.

**Figure 3.**
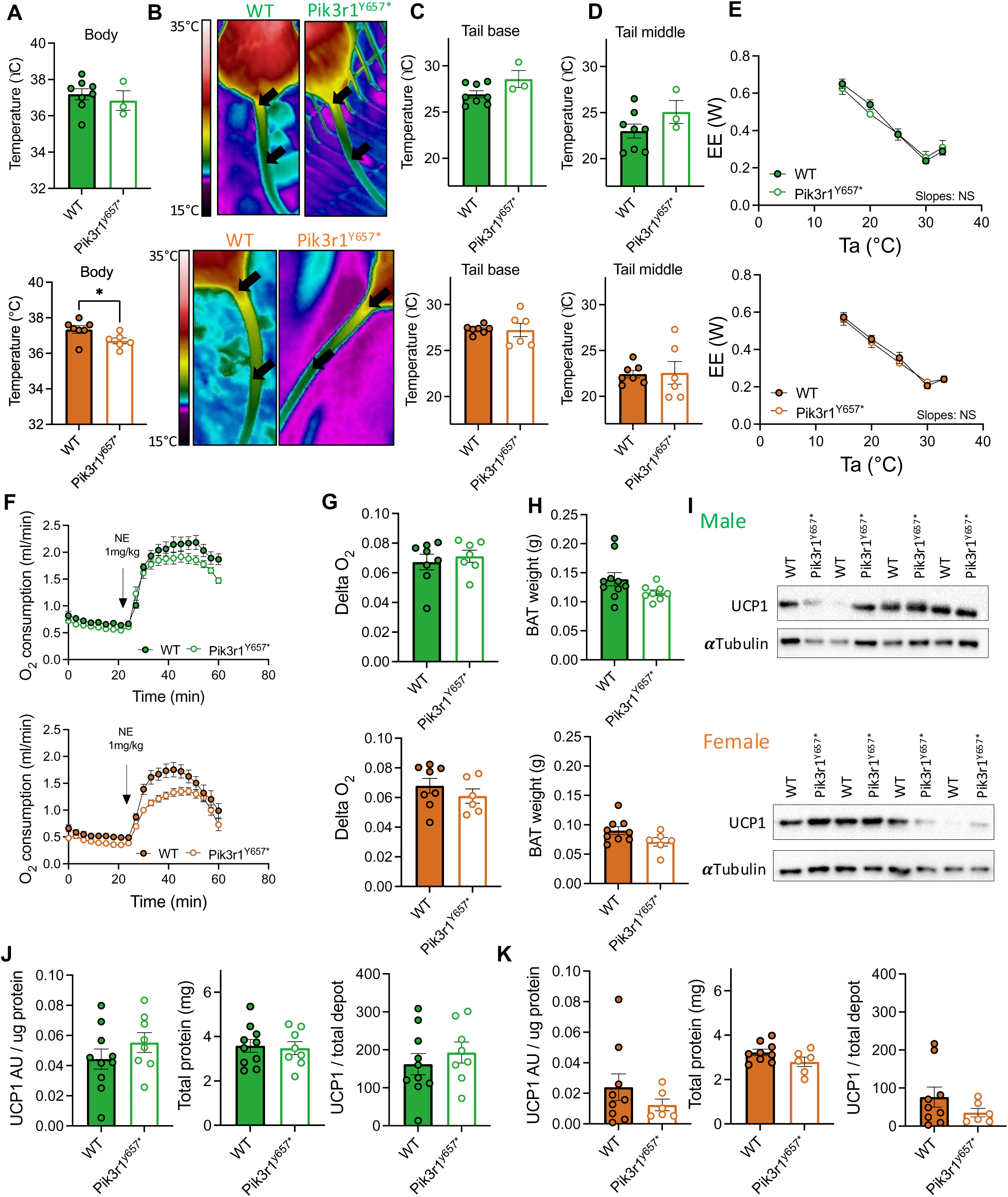
Tail vein heat dissipation, insulation, and Ucp1-dependent thermogenic capacity in *Pik3r1^Y657*^* mice at 21°C. Male (green; top in paired images) and female (orange; below in paired images) WT and *Pik3r1^Y657*^* mice were fed a 45% HFD and kept at 21°C throughout the study. (A) Body temperature measured from the inner ear at 14 weeks old. (B) Representative thermal images at 14 weeks old. Black arrows indicate tail areas used for quantification of temperature. (C) Quantification of tail base temperature. (D) Quantification of temperature at the middle of the tail. (E) Energy expenditure in relationship to ambient temperature (Ta). (F) Norepinephrine (NE)-induced oxygen consumption in pentobarbital-anaesthetized mice. (G) Quantification of (F): Ucp1-dependent thermogenic capacity = peak NE-induced O_2_ consumption – basal O_2_ consumption. (H) Total BAT weight. (I) Representative immunoblots showing Ucp1 protein in BAT homogenates with αTubulin as a loading control. (J,K) Quantification of (I) showing Ucp1 per microgram protein, total protein per depot, and total Ucp1 per depot. All data are represented as mean ± SEM, error bars may be too small to be visible; Male WT N=8, Male *Pik3r1^Y657*^* N=3, Female WT N=7, Female *Pik3r1^Y657*^* N=6 (A, C, D). Male WT N=6, Male *Pik3r1^Y657*^* N=3, Female WT N=6, Female *Pik3r1^Y657*^*N=7 (E). Male WT N=10, Male *Pik3r1^Y657*^* N=8, Female WT N=9, Female *Pik3r1^Y657*^* N=6 (F-J). NS = non-significant, * P<0.05, ** P<0.01. Student’s t test (A, C, D, G-K); Linear regression on data points from 15°C to 30°C, slope and Y-intercept comparison between genotypes (E).

Thermal insulation is also a determinant of cold sensitivity and thus the amount of energy used to defend body temperature. For mice, insulation is mainly determined by fur and skin thickness [31,32], both factors influenced by the PI3K/Akt signaling pathway (e.g. [33–35]). To determine whether insulation was decreased in *Pik3r1^Y657*^* mice, we performed a modified Scholander experiment [27]. We found that for both WT and *Pik3r1^Y657*^* mice, 30°C was the lower critical temperature for the thermoneutral zone (Fig. 3E). When ambient temperature dropped below 30°C, both genotypes of both sexes increased energy expenditure to defend their body temperature, as expected. We found that the incremental increases in energy expenditure per degree reduction in ambient temperature (Fig. 3E, slopes) were identical between genotypes, indicating no difference in insulation between genotypes or sexes.

Finally, we determined total Ucp1-dependent non-shivering thermogenic capacity. Although the similarities in thermoregulatory phenotype between *Pik3r1^Y657*^* and WT mice reported here (Figs. 3A-E) render BAT hyperactivation unlikely as a response to increased heat loss, reduced PI3K activity has been suggested to affect BAT *Ucp1* expression directly, and thus could increase BAT activity in a cell- or tissue-autonomous manner [12–14,17]. To determine total Ucp1-dependent thermogenic capacity in *Pik3r1^Y657*^*and WT mice, we uncoupled all Ucp1 present in pentobarbital-sedated mice by NE injection and recorded the resulting increase in O_2_ consumption (Fig. 3F). Mice of both sexes and genotypes increased O_2_ consumption after NE injection, but we found no difference in Ucp1-dependent thermogenic capacity (i.e. peak NE-induced O_2_ consumption – basal O_2_ consumption (Fig. 3G). To confirm this finding, we dissected interscapular BAT and inguinal WAT and determined Ucp1 protein levels. No differences were found between genotypes in BAT tissue weight or total protein content (Figs. 3H, 3J&K left panels), and indeed no differences were found in BAT Ucp1 protein levels, either when expressed per ug protein or calculated per total depot (Figs. 3J, K). No detectable levels of Ucp1 were found in ingWAT, thus also confirming no changes in WAT browning (Suppl. Fig. 1).

### Increased energy expenditure persists in male, but not female, Pik3r1^Y657*^ mice at thermoneutrality

In the studies above (Figs. 1-3), we have shown that, despite increased energy expenditure, there is no evidence that *Pik3r1^Y657*^* mice have altered thermoregulation or BAT activation. However, these studies were performed in mice housed at 21°C, below murine thermoneutrality, which requires mice to double their energy expenditure to defend their body temperature (Fig. 3E). This makes it difficult to interpret and extrapolate results from these murine studies to humans, since humans live most of their lives at thermoneutrality. To enhance the potential for extrapolation of our results to human SHORT syndrome, we thus next studied the mice in a ‘thermally humanized’ environment. From 10 weeks of age, WT and *Pik3r1^Y657*^*mice were housed at thermoneutrality (30°C), and after 2-weeks’ acclimatisation, received HFD for 3 weeks.

Being housed at thermoneutrality did not alter the gross phenotype of the mice, as both male and female *Pik3r1^Y657*^* mice had reduced body weight and length compared to their WT counterparts (Figs. 4A, B). Tibia length was also reduced in male *Pik3r1^Y657*^* mice, but not in females (Fig. 4C). As for mice housed at 21°C, male and female *Pik3r1^Y657*^* mice housed at 30°C had much reduced lean mass compared to WT mice throughout the study (Fig. 4D), with the pattern of fat mass accumulation upon HFD-feeding also comparable between temperatures (Figs. 1F versus 4E).

**Figure 4.**
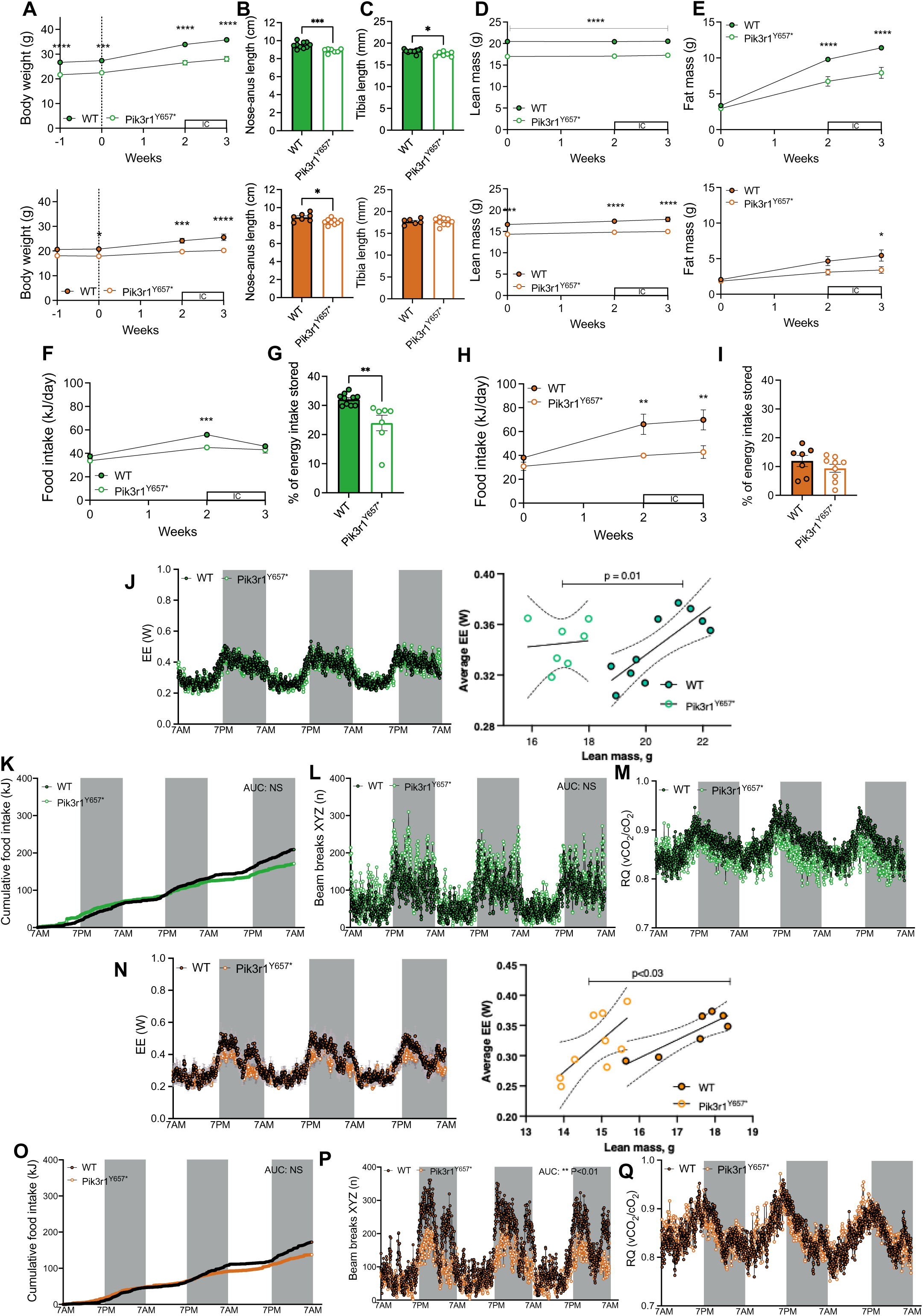
Energy metabolism of *Pik3r1^Y657*^* mice at 30°C. Male (green) and female (orange) WT and *Pik3r1^Y657*^* mice were moved from 21°C to 30°C ambient temperature at 10 weeks’ old, and then fed a 45% HFD from 12 weeks’ old (t=0, indicated by dotted lines), being kept at 30°C for the remainder of the study. Boxes with ‘IC’ represent periods in the indirect calorimetry system. Grey boxes represent dark phases. (A) Body weight over time. (B) Body length (nose-anus) at 15 weeks’ old. (C) Tibia length at 15 weeks’ old. (D) Lean mass over time. (E) Fat mass over time. (F, H) Food intake. (G, I) Metabolic efficiency calculated between 12-14 weeks old. (J,N) 72h energy expenditure. Raw traces are shown with adjacent linear regression plots of the last 24 hours’ energy expenditure against lean mass. Regression lines and 95% confidence interval are shown, with p value relating to testing of differences in intercept (no difference in gradients) (K,O) Cumulative food intake calculated in kJ. (L,P) Sum of X, Y and Z beam breaks per time point. (M,Q) Respiratory quotient. All data are represented as mean ± SEM (shaded areas in J-M); error bars may be too small to be visible; Male WT N=10, Male *Pik3r1^Y657*^* N=7, Female WT N=7, Female *Pik3r1^Y657*^* N=9. NS = non-significant, * P<0.05, ** p<0.01, *** p<0.001 and **** p<0.0001. Two-way ANOVA with Šídák’s multiple comparisons test, significance per timepoint between genotypes (A, D-F, H); Student’s t test (B, C, G, I), AUC were determined per animal and averaged per genotype, compared by Student’s t test (J-L).

Thermoneutrality abolished the baseline difference in food intake between male WT and *Pik3r1^Y657*^* mice suggested at 21°C (Fig. 4F), but reduced metabolic efficiency (Fig. 4G), and increased energy expenditure (Fig. 4J) persisted. Increased energy expenditure of male *Pik3r1^Y657*^*mice at thermoneutrality could not be explained by increased food intake or physical activity (Figs. 4K, L). Respiratory quotient was lower in the dark phase than in the light, suggesting increased reliance on fatty acid oxidation (M). In females, no baseline difference in food intake was found between WT and *Pik3r1^Y657*^* mice at 30°C (Fig. 4H, K). However, in contrast to the males, the reduced metabolic efficiency seen in female *Pik3r1^Y657*^* mice at 21°C (Fig. 1H) was not evident at 30°C (Fig. 4I), indicating that thermal stress may influence the *Pik3r1^Y657*^* energetic phenotype in females. Female *Pik3r1^Y657*^* mice at 30°C did, however, have increased energy expenditure compared to WT mice (Fig. 4N), despite no change in food intake (Fig 4O) and a decrease in physical activity (Fig. 4P). No difference in respiratory quotient was seen.

### No change in Ucp1-dependent thermogenic capacity at thermoneutrality

Although we did not find an increase in BAT thermogenic capacity that could explain the increased energy expenditure in *Pik3r1^Y657*^* mice at 21°C, thermal stress might complicate interpretation of these results. At 21°C, BAT activity is mainly determined by ongoing sympathetic output to the tissue, and such continuous activation may mask direct effects of impaired PI3K signaling on BAT. Thus, we finally determined Ucp1-dependent thermogenic capacity in WT and *Pik3r1^Y657*^* mice housed at thermoneutrality.

As shown in Fig. 5A, the increase in O_2_ consumption after NE injection was much lower in both male and female WT and *Pik3r1^Y657*^* mice housed at 30°C than in mice housed at 21°C (Fig. 3F), reflecting the expected involution of BAT in the absence of hypothermic stress. However, no difference in NE-induced O_2_ consumption was again seen between genotypes (Figs. 5A, B). This was confirmed by western blot analyses (Figs. 5C-E, Suppl. Fig. 2), again indicating no increase in Ucp1-dependent thermogenesis in *Pik3r1^Y657*^*mice.

**Figure 5.**
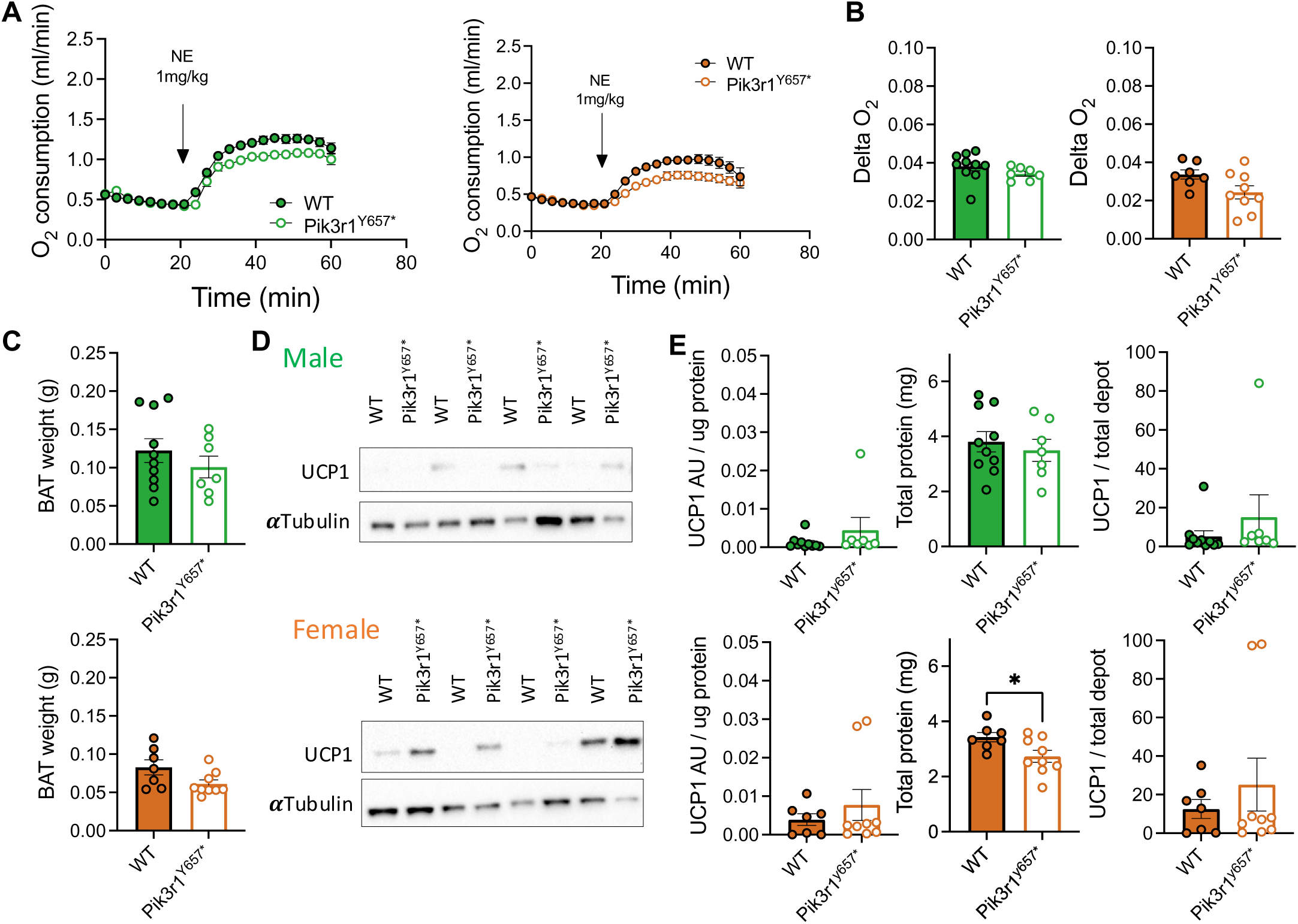
No BAT hyperactivation in *Pik3r1^Y657*^* mice at 30°C. Male (green; left or top in paired images) and female (orange; right or bottom in paired images) WT and *Pik3r1^Y657*^* mice were moved from 21°C to 30°C ambient temperature at 10 weeks’ old, and then fed a 45% HFD from 12 weeks’ old. (A) Norepinephrine (NE, 1mg/kg)-induced oxygen consumption in pentobarbital-anaesthetized mice. (B) Quantification of (A): Ucp1-dependent thermogenic capacity = peak NE-induced O_2_ consumption – basal O_2_ consumption. (C) Total BAT weight. (D) Representative immunoblots showing Ucp1 protein in BAT homogenates with αTubulin as a loading control. (E) Quantification of (D) showing Ucp1 per microgram protein, total protein per depot, and total Ucp1 per depot. All data are represented as mean ± SEM; error bars may be too small to be visible; Male WT N=10, Male *Pik3r1^Y657*^* N=7, Female WT N=7, Female *Pik3r1^Y657*^* N=9. * P<0.05. Two-way ANOVA with Šídák’s multiple comparisons test, significance per timepoint between genotypes (A); Student’s t test (B-E).

## Discussion

Fatty liver and metabolic dyslipidaemia are common, major complications of idiopathic, genetic and acquired forms of IR. However, severe IR caused by defects in the insulin receptor or PI3K is not associated with these complications [5,36]. Studying proximal insulin signalling defects may thus provide novel, important insights into approaches for mitigating IR-associated complications.

We previously showed that *Pik3r1^Y657*^*mice phenocopy human SHORT syndrome, featuring severe IR and reduced adiposity, but neither fatty liver nor dyslipidemia [6]. We now confirm the increase in energy expenditure that we reported previously in male *Pik3r1^Y657*^* mice at 21°C in both males and females, when their smaller lean mass is accounted for [6]. This finding is consistent with growing evidence that inhibiting PI3K signaling pharmacologically or genetically stimulates energy expenditure in rodents and primates [12–19]. Although SHORT syndrome is often described as featuring lipodystrophy, our interpretation is that the reduced adipose tissue with healthy lipid phenotype and preserved serum adiponectin concentration[5] is a result of increased energy expenditure. Lipodystrophy usually features adipose failure, which we have previously failed to demonstrate in *Pik3r1^Y657*^*mice, but we do consistently find increased energy expenditure. This remains to be tested in humans, however.

To investigate the mechanism of increased energy expenditure in *Pik3r1^Y657*^* mice, we tested different components of energy expenditure, ruling out increased obligatory diet-induced thermogenesis or physical activity as explanations. We also found no indication of increased Ucp1-dependent thermogenesis, whether secondary to other changes in thermoregulation, or tissue-autonomous. This is surprising, as in prior studies in an array of rodent models of impaired PI3K signalling, increased BAT thermogenesis was suggested [12–14,17,18]. This was not the case in all models, however, as no involvement of BAT [37] and even a downregulation of BAT activity has also been reported [21]. The discrepancy between the results reported here and the literature may reflect the measurements used. Several studies have reported increased glucose uptake into BAT [13,14] and upregulation of *Ucp1* gene expression in BAT and ingWAT [12–14,17]. However, BAT glucose uptake is disconnected from the amount of Ucp1 in, and thus thermogenic capacity of, BAT [38]. Moreover, the use of *Ucp1* mRNA levels as a proxy for BAT recruitment may be inadequate, as Ucp1 only contributes to energy expenditure when uncoupled upon adrenergic stimulation [39]. We have here directly assessed total thermogenic capacity by injecting NE into anaesthetized WT and *Pik3r1^Y657*^*mice and quantifying the increase in O_2_ consumption. Similar measurements have been reported in one other study [13], but increased basal O_2_ consumption due to impaired PI3K signalling was unaccounted for in that analysis. We here expressed Ucp1-dependent thermogenic capacity as difference between peak NE-induced and baseline O_2_ consumption, and found no difference between genotypes.

Nearly all studies investigating mice with impaired PI3K signaling have been performed in mice housed at 21°C, i.e. below murine thermoneutrality [24]. We show that without thermal stress, the energy expenditure increase remains in both male and female *Pik3r1^Y657*^* mice, but is mitigated in female mice by decreased physical activity. We further report no difference in whole body Ucp1-dependent thermogenic capacity between male and female *Pik3r1^Y657*^* mice at 30°C. This suggests that, contrary to previous suggestions [13,14], impaired PI3K signalling in itself does not increase sympathetic output to thermogenic adipose tissue, nor increase *Ucp1* expression in a cell autonomous manner.

We provide conclusive evidence that the increased energy expenditure in a SHORT syndrome model is not caused by changes in physical activity, diet-induced thermogenesis, thermoregulation, or Ucp1-mediated energy dissipation in BAT. Previous study of *Pik3r1^Y657*^* mice also ruled out increased energy loss via nutritional malabsorption [6]. This leaves the question of the mechanism underlying increased energy expenditure in *Pik3r1^Y657*^* mice unresolved.

One explanation put forward for increased energy expenditure induced by impaired PI3K signaling is an overall increase in mitochondrial oxidative phosphorylation, potentially translating into an increased basal metabolic rate (BMR) [13,14,16,37]. Another non mutually exclusive possibility is increased metabolic futile cycling. Although Ucp1-mediated proton flux is the best studied futile cycle, several others are known, [40–44]. There is keen interest in exploiting such cycles to combat obesity and metabolic disease [45]. Metabolic fluxes have not yet been assessed in *Pik3r1^Y657*^* mice, however metabolic assessments in fed and fasting states have suggested increased net fatty acid oxidation in the fed state with altered profiles of plasma amino acids. Finally, our previously reported observation that *Pik3r1^Y657*^* mice have larger hearts than WT mice when corrected for body mass [6] may also hold clues. The heart accounts for around 10% of basal metabolic rate in healthy humans [46], variations in its size have been suggested to contribute to alterations in BMR [47,48], and in some circumstances the heart can influence energy expenditure in remote tissues through circulating mediators [49,50]. Whether the enlarged hearts in *Pik3r1^Y657*^* mice consume more energy themselves, and whether the enlarged heart influences other tissues in this case remains to be determined.

## Conclusion

Male and female *Pik3r1^Y657*^* mice at 21°C and 30°C, show increased lean mass-adjusted energy expenditure that we suggest protects from lipotoxicity despite IR. The increased energy expenditure cannot be explained by changes in physical activity, diet-induced thermogenesis, or increased Ucp1-dependent thermogenesis, whether secondary to other changes in thermoregulation, or tissue-autonomous. More research is needed into the mechanism underlying increased energy expenditure in *Pik3r1^Y657*^* mice.

## CRediT authorship contribution statement

**IL:** conceptualization, methodology, formal analysis, investigation, writing - original draft, writing - review & editing, visualization, funding acquisition. **AO:** methodology, investigation, writing - review & editing. **EM:** investigation, writing - review & editing. **TB:** resources, writing - review & editing. **RKS:** conceptualization, resources, writing - original draft, writing - review & editing, supervision, funding acquisition.

## Acknowledgements

We thank Jan Nedergaard and Barbara Cannon for kindly gifting the Ucp1 antibody and for providing feedback on methodology. We thank Roland Stimson for providing access to the FLIR thermal camera, and Nik Morton for access to the TD-NMR machine at the University of Edinburgh. We are grateful for the excellent technical assistance at the Biological Research Facility at the University of Edinburgh and at the Experimental Core Facility at Stockholm University.

---

**Supplementary Figure 1, Related to Fig. 3.**
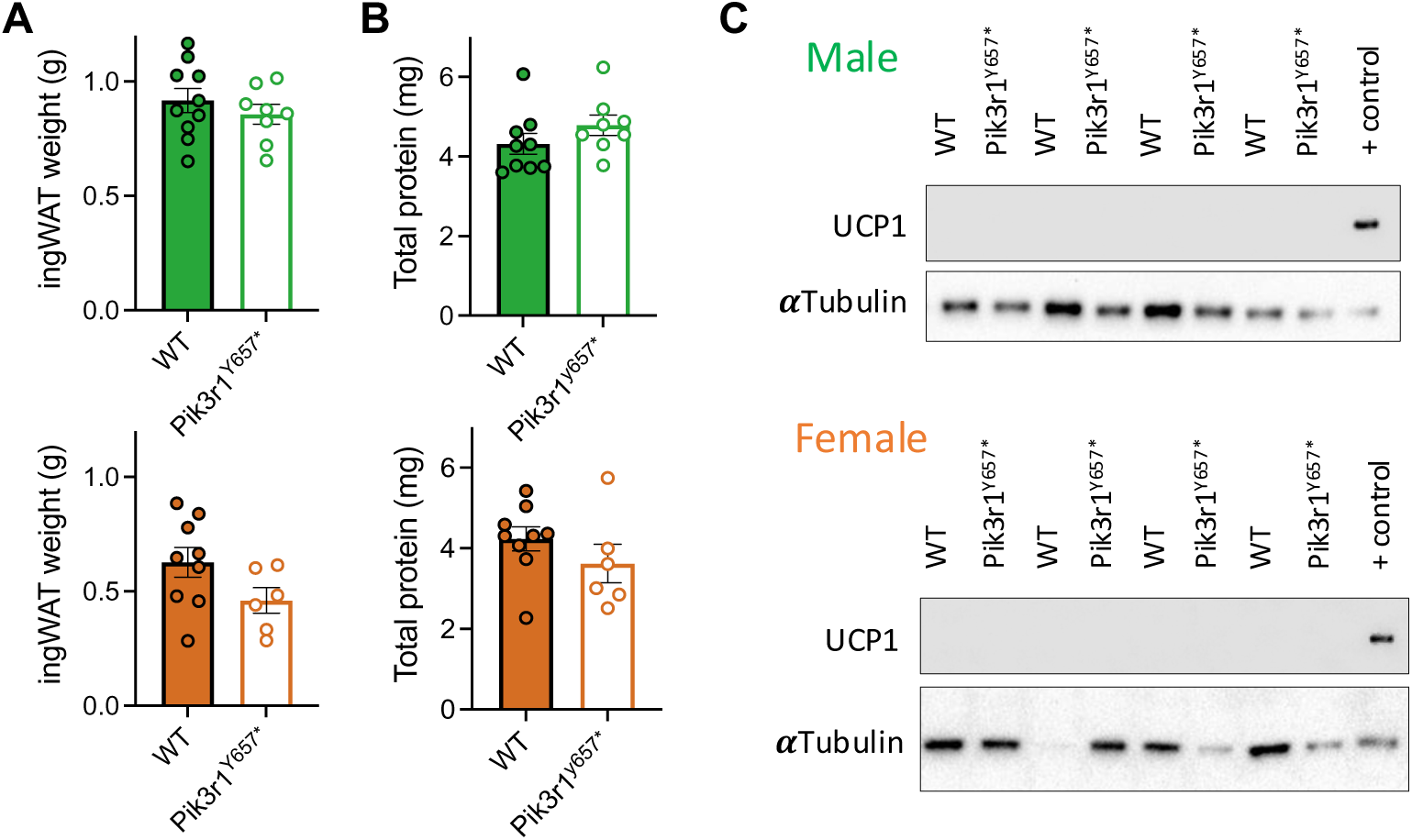
Ucp1 protein amount in inguinal WAT at 21 °C. Male (green) and female (orange) WT and Pik3r1^Y657*^ mice were fed a 45% HFD and kept at 21 °C for the duration of the study. (A) Total ingWAT weight. (B) Total protein per depot. (C) Representative immunoblots showing Ucp1 protein in BAT homogenates with αTubulin as a loading control. All data are represented as mean ± SEM; Male WT N = 10, Male Pik3r1^Y657*^ N = 8, Female WT N = 9, Female Pik3r1^Y657*^ N = 6. Student’s t test (all).

**Supplementary Fig. 2, Related to Fig 5.**
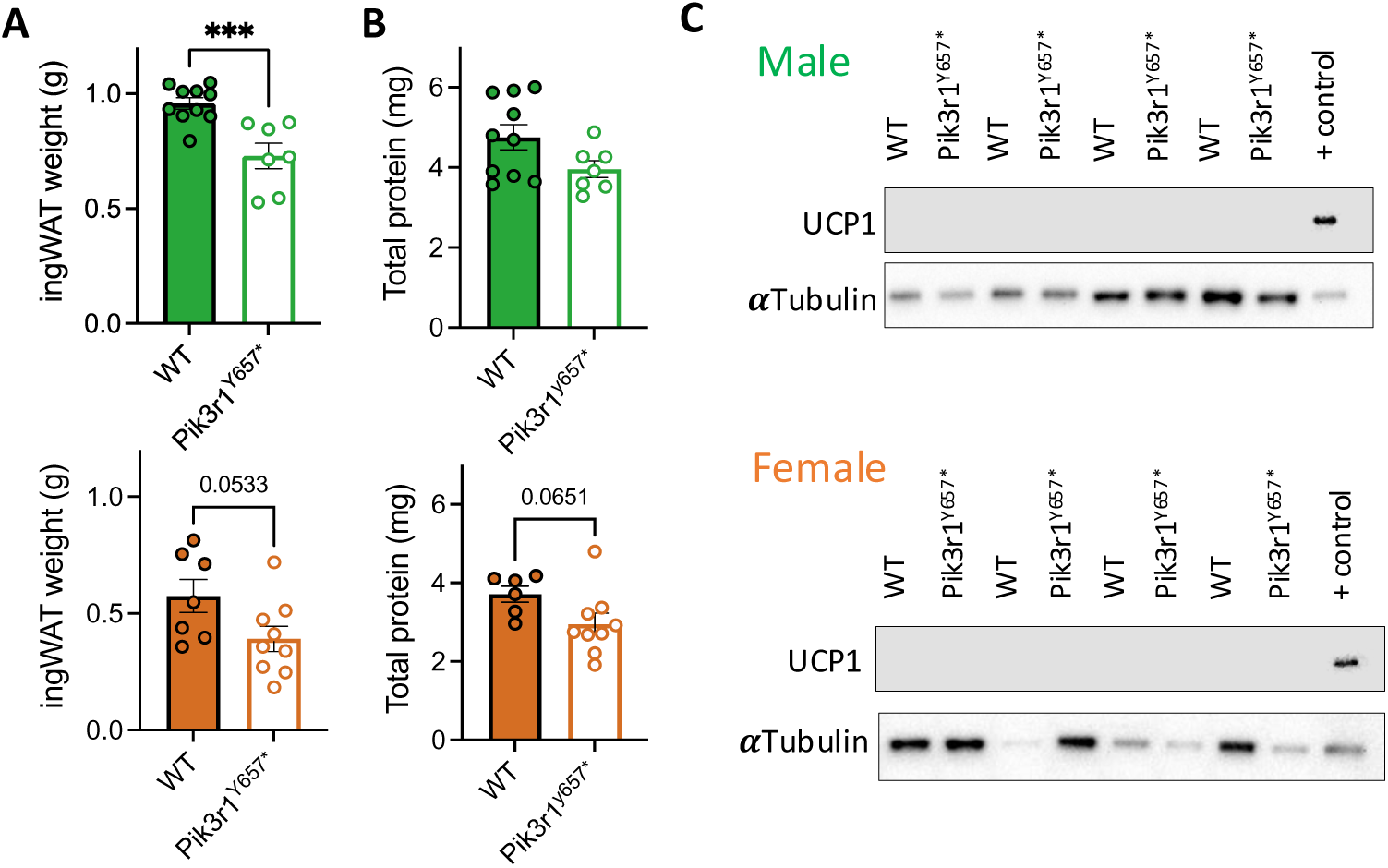
Ucp1 protein amount in inguinal WAT at 30 °C. Male (green) and female (orange) WT and Pik3r1^Y657*^ mice were fed a 45% HFD and kept at 30 °C for the duration of the study. (A) Total ingWAT weight. (B) Total protein per depot. (C) Representative immunoblots showing Ucp1 protein in BAT homogenates with αTubulin as a loading control. All data are represented as mean ± SEM; Male WT N = 10, Male Pik3r1^Y657*^ N = 7, Female WT N = 7, Female Pik3r1^Y657*^ N = 9. *** p<0.001, Student’s t test (all).

